# Rapid and accurate identification of *Escherichia coli* STEC O157:H7 by mass spectrometry, artificial intelligence and detection of specific biomarkers peaks

**DOI:** 10.1101/2022.03.31.486435

**Authors:** Manfredi Eduardo, Rocca María Florencia, Zintgraff Jonathan, Irazu Lucía, Miliwebsky Elizabeth, Carbonari Carolina, Deza Natalia, Prieto Monica, Chinen Isabel

**Affiliations:** Servicio Fisiopatogenia, Instituto Nacional de Enfermedades Infecciosas (INEI) – Administración Nacional de Laboratorios e Institutos de Salud (ANLIS) “Dr. Carlos G. Malbrán”; Instituto Nacional de Enfermedades Infecciosas (INEI) – Administración Nacional de Laboratorios e Institutos de Salud (ANLIS) “Dr. Carlos G. Malbrán”; Red Nacional de Espectrometría de Masas aplicada a la Microbiología Clínica (ReNaEM Argentina), Argentina

**Author notes:** **Corresponding author at:** *Instituto Nacional de Enfermedades Infecciosas (INEI) – Administración Nacional de Laboratorios e Institutos de Salud (ANLIS) “Dr. Carlos G. Malbrán* **E-mail addresses:** (E.M), (M.F.R), (J.Z). These authors contributed equally to this work (Manfredi Eduardo, Rocca María Florencia, Zintgraff Jonathan).

## Abstract

The different pathotypes of *Escherichia* can produce a large number of human diseases. Surveillance becomes complex since their differentiation are not easy.

Particularly, the detection of Shiga toxin-producing *Escherichia coli* (STEC) serotype O157:H7 consists of stool culture of a diarrheal sample in enrichment and/or selective media, identification of presumptive colonies and confirmation by Multiplex PCR technique for the genotypic characterization of serogroup O157 and Shiga toxins (*stx*1 and *stx*2), in addition to the traditional biochemical identification.

All of these procedures are laborious, require a certain level of training, are time consuming and expensive. Among the currently most widely used methodologies, MALDI-TOF MS mass spectrometry (matrix-assisted laser desorption/ionization with time-of-flight mass detection), allows a quick and easy way to obtain a protein spectrum of a microorganism, not only in order to identify the genus and species, but also the discovery of potential biomarker peaks of a certain characteristic. In the present work, the information obtained from 60 clinical isolates was used to detect peptide fingerprints of STEC O157:H7 and other diarrheagenic *E. coli*. The differences found in the protein profiles of the different pathotypes established the foundations for the development and evaluation of classification models through automated training.

The application of the Biomarkers in combination with the predictive models on a new set of samples (n=142), achieved 99.3% of correct classifications, allowing the distinction between STEC O157:H7 isolates from the other diarrheal *Escherichia coli*.

Therefore, given that STEC O157:H7 is the main causal agent of haemolytic uremic syndrome and based on the performance values obtained in the present work (Sensitivity=98.5% and Specificity=100%), this development could be a useful tool for diagnosis of the disease in clinical microbiology laboratories.

## INTRODUCTION

The contribution of *Escherichia coli* to human intestinal disease may be largely uncharacterized, because many types of pathogenic *E. coli* are not routinely tested in clinical microbiology laboratories.

Shiga toxin-producing *Escherichia coli* (STEC) is associated with outbreaks that causes diarrhea, hemorrhagic colitis, and hemolytic-uremic syndrome (HUS) in humans. It is part of the diarrheagenic *E. coli* (DEC) group, which also includes: enteropathogenic *E. coli* (EPEC), enterotoxigenic *E. coli* (ETEC), enteroaggregative *E. coli* (EAEC), enteroinvasive *E. coli* (EIEC) and diffusely adherent *E. coli* (DAEC). Although, there are more than 150 serotypes **[1]** that share the same pathogenic potential, O157:H7 is the most frequent. In particular, the detection of STEC O157:H7 consists of the culture of the faecal sample in enrichment and/or selective media such as MacConkey agar with sorbitol for the identification of presumptive non-fermenting sorbitol colonies and confirmation by Multiplex PCR for the genotypic characterization of serogroup O157 and Shiga toxins (*stx*1 and *stx*2), in addition to subsequent traditional biochemical identification.

However, all this methodological complexity is very difficult to implement in a traditional laboratory **[2]**.

On the other hand, mass spectrometry (MS), specifically MALDI-TOF MS (matrix-assisted laser desorption/ionization with time-of-flight mass detection), provides a simple, rapid, robust, and low-cost microbial identification. MALDI-TOF MS is a technique based on the analysis of protein spectra containing peaks with an exactly determinable mass-charge ratio (m/z) generated by the impact of a laser on a previously crystallized isolate with an organic matrix. In recent years, MS has acquired great importance in the identification of pathogens that are clinically relevant in public health **[3-5]**. However, the potential of this methodology combined with machine learning algorithms for the detection of profiles in a wide variety of samples and its use as a screening technique is expanding, due to its low-cost and high performance **[6]**.

In this study, we wanted to verify the usefulness of MS to rapidly identify O157:H7 STEC from pathotypes other than diarrheagenic *E. coli*; then, we proposed to detect and analyse peaks in the spectra generated by MALDI-TOF MS to find possible biomarkers and thus establish differential patterns between a wide variety of *E. coli* strains.

## MATERIALS AND METHODS

### Isolates collection

The spectra obtained from 60 isolates corresponding to four different DEC categories were used for the development of predictive models and the detection of biomarkers: EPEC (n=15), ETEC (n=15), STEC NON O157 (n=20) and STEC O157:H7 (n=10). The detail of the isolates can be found in **Table S1** in Supplementary Material.

For the final validation, we used 142 different isolates of: STEC O157:H7 (n=65), non-toxigenic *E*.*coli* O157 (n=13), STEC NON O157 (n=17), ETEC (n= 11), EPEC (n=12), EAEC (n=15), EIEC (n=7), and *E. coli* without virulence factors (n=2). All of them were obtained mainly from faecal samples from different health institutions in our country and food samples subsequently referred to the National Reference Laboratory-Servicio de Fisiopatogenia INEI-ANLIS “Dr. Carlos G. Malbrán”-for confirmation and characterization of specific virulence factors **[7]**.

### Acquisition of spectra

The Microflex LT mass spectrometry equipment (Bruker Daltonics) was used to obtain the protein spectra from *E. coli* isolates. Subsequently, each isolate was spotted in quadruplicate in the wells of the steel plate provided by the manufacturer using the direct method and crystallized by adding 1 ul of HCCA matrix (α-Cyano-4-hydroxycinnamic acid in 50% of acetonitrile and 2.5% trifluoroacetic acid). After a few minutes of drying, the plate was introduced into the equipment and once the vacuum was reached in the Flex Control v3.4 software, the spectra were acquired in the linear positive mode, with 30-40% laser power. and in a mass range of 2 to 20 kDa. Each well was read twice, so eight individual spectra were obtained for each isolate, thus minimizing the variability of the technique.

The external calibrator provided by the manufacturer, BTS (Bruker Test Standard), was used prior to each run.

The 142 isolates used in the subsequent validation set were processed in the same way by direct method, but each isolate was spotted in duplicate and read only once, simulating the routine procedure of a microbiology laboratory.

### MALDI-TOF analysis

All isolates were identified using the MALDI Biotyper RTC software. According to the manufacturer’s recommendations, the identification is considered reliable at the species level when the score values greater than 2.0 are obtained. When the score values are between 1.7 and 1.99, it is considered reliable identification at the genus level; and it is considered ‘No Identification’ when the value of the score is ≤ 1.69 **[8]**.

### Bioinformatic analysis

To perform data analysis, ClinPro Tools software (version 3.0, Bruker Daltonik GmbH, Bremen, Germany) and Flex Analysis v3.4 software (Bruker Daltonics, Bremen, Germany) were used.

### Data pre-processing

In order to take advantage of the greatest amount of information contained in the spectra, the following data pre-processing steps were performed: baseline correction (top hat 10% of minimum width of the baseline), smoothing and calibration excluding null spectra or out of range, according to the literature **[9-11]**.

### Peak Selection

The exploration of the potential biomarkers was performed on the protein profiles generated from the 60 isolates that were part of the initial training group and which were also, used to create the classification models.

### Flex Analysis v3.4 Software

All spectra were exported as mzXML files using CompasXport CXP3.0.5 and a series of analyses were performed according to standard Bruker setup. Spectrum quality criteria for overall aspect and intensity were checked. Next, visually identifiable biomarker peaks were explored in the different views offered by the program. The mass list was exported to Excel (Microsoft, Redmond, WA) to analyse possible biomarker peaks. Values of “1” or “0” (data binarization) were assigned to the presence or absence of a peak within the tolerance interval (+/-10Da). Based on this analysis, groups of potentially useful peaks for the diagnosis of STEC O157:H7 were found.

### ClinProTools v3.0 Software

The spectra of STEC O157:H7 were assigned as class 1 and the rest of the DEC isolates were class 2. Biomarker peaks were automatically identified by class comparison using the function “Peak Statistic Table”.

To select the characteristic peaks of the two classes, the following statistical tests were used: the t test/analysis of variance ANOVA (PTTA), Wilcoxon or Kruskal–Wallis test (W/KW), and Anderson–Darling test (AD). A p value of 0.05 was established as the cut-off point **[12]**:

- if p is <0.05 in the AD test, a characteristic peak is selected if the corresponding p-value in the W/KW test is also <0.05.
- if p is 0.05 in the AD test, then a characteristic peak is selected if the corresponding p value in ANOVA is also <0.05 **[13]**.

The discriminative power for each biomarker was further described by receiver operating characteristic (ROC) and area under the curve (AUC) analysis.

ROC curve indicates the relationship of the true-positive rate (TPR) and the false-positive rate (FPR). The area under the ROC curve is equal to the probability that a biomarker sorts a randomly selected positive sample higher than a randomly selected negative one.

### Principal Component Analysis (PCA)

To explore and compare spectra in multidimensional space and in order to evaluate the possible distributions or clusters on the isolates of both classes, a first exploratory and unsupervised analysis was performed of all 60 samples.

### Classification models

Supervised classification models were performed using the following algorithms provided by ClinPro Tools software: Supervised Neural Network (SNN), optimized genetic algorithm combined with k-nearest neighbour classification (GA/ kNN) and a quickclassifier (QC).

For each model, the following parameters were calculated: Recognition Capability (CR) and Cross Validation Percentage (VC), both of which are indicators of the theoretical behaviour that the model will have in future classifications.

### Selection of isolates for final validation

To evaluate the robustness of the models created, an independent set of isolates different from those used to developed the algorithms was selected (N=142). For each isolate, a spectrum was presented to the selected classification model. The software then returned a result that was compared to current reference techniques.

### Statistical analysis

The parameters evaluated were: sensitivity, specificity, positive predictive value, negative predictive value **[14]**. Besides, the CLSI guide, EP12-A2, was used to compare methods that report results qualitatively. When the comparison is made with a method that is not considered a reference, the degree of similarity between the methods is measured through the percentage of negative agreement and the percentage of positive agreement. The diagnostic parameters of the methods are then compared to determine if the difference between the two of them is statistically significant.

## RESULTS

### Confirmation at the genus-species level

All isolates were identified at the species level as *Escherichia coli* with a score value greater than 2.0, in agreement with the result of the reference techniques.

### Unsupervised Analysis

First, a Principal Component Analysis (PCA) was performed for the 60 isolates used as a training set. In this way it was possible to reduce all the information contained in the MALDI-TOF spectra in a few new variables.

This allows us to graphically represent all the spectra together in three and two dimensions on the Score plot. (Figure 1) The complete list of significant peaks found in ClinPro Tools can be found in **Supplementary Material S2**.

**Figure 1.**
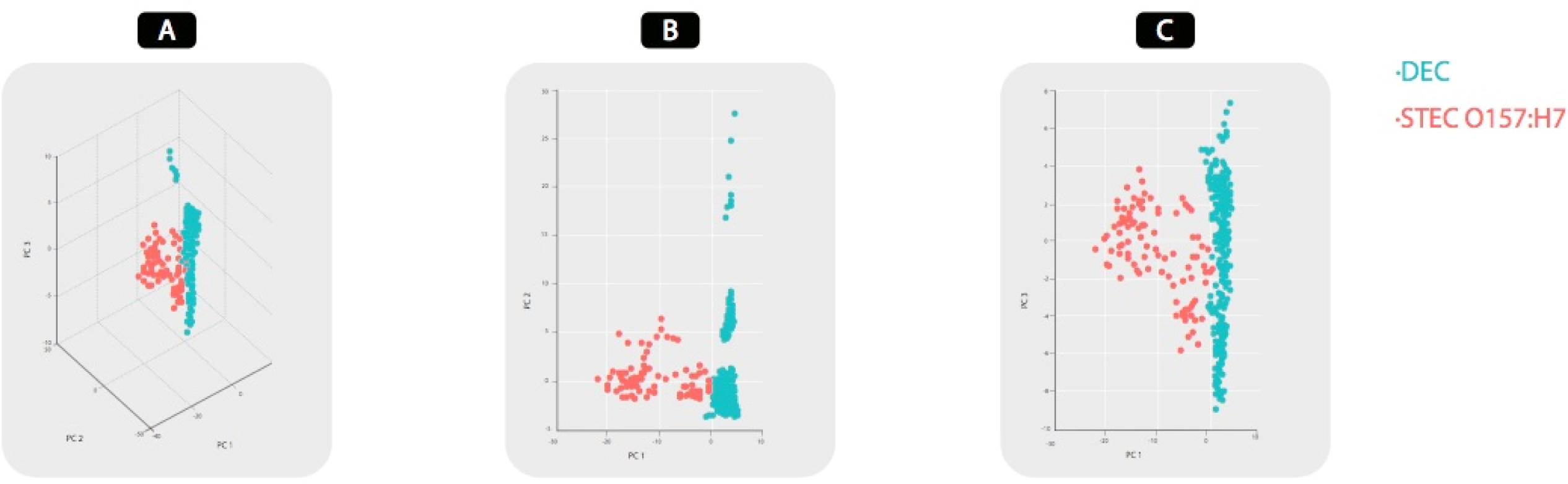
PCA plots results. Data originated from the external MATLAB software tool integrated into ClinPro Tools.

Graph A shows the three components plotted simultaneously in three dimensions, while in graphs B and C shows PC1 versus PC2 and PC1 versus PC3 respectively.

### Supervised Analysis

Subsequently, a supervised multivariate analysis was performed with the additional information of each isolate to define each class:

**CLASS 1: STEC O157:H7**

**CLASS 2: NON STEC O157 (other DEC)**

Figure 2-A shows the two-dimensional distribution plot of all the spectra of each class based on the two best peaks obtained for their classification; which correspond to the 9137.26 Da peak and the 9227.11 Da peak. The peak number and its m/z values are shown on the x and y axes respectively, while the ellipses represent the 95% confidence interval. On the other hand, Figure 2-B shows the ROC curves of the two selected peaks. The area under the curve (AUC) represents the discriminatory potential of each biomarker peak.

**Figure 2.**
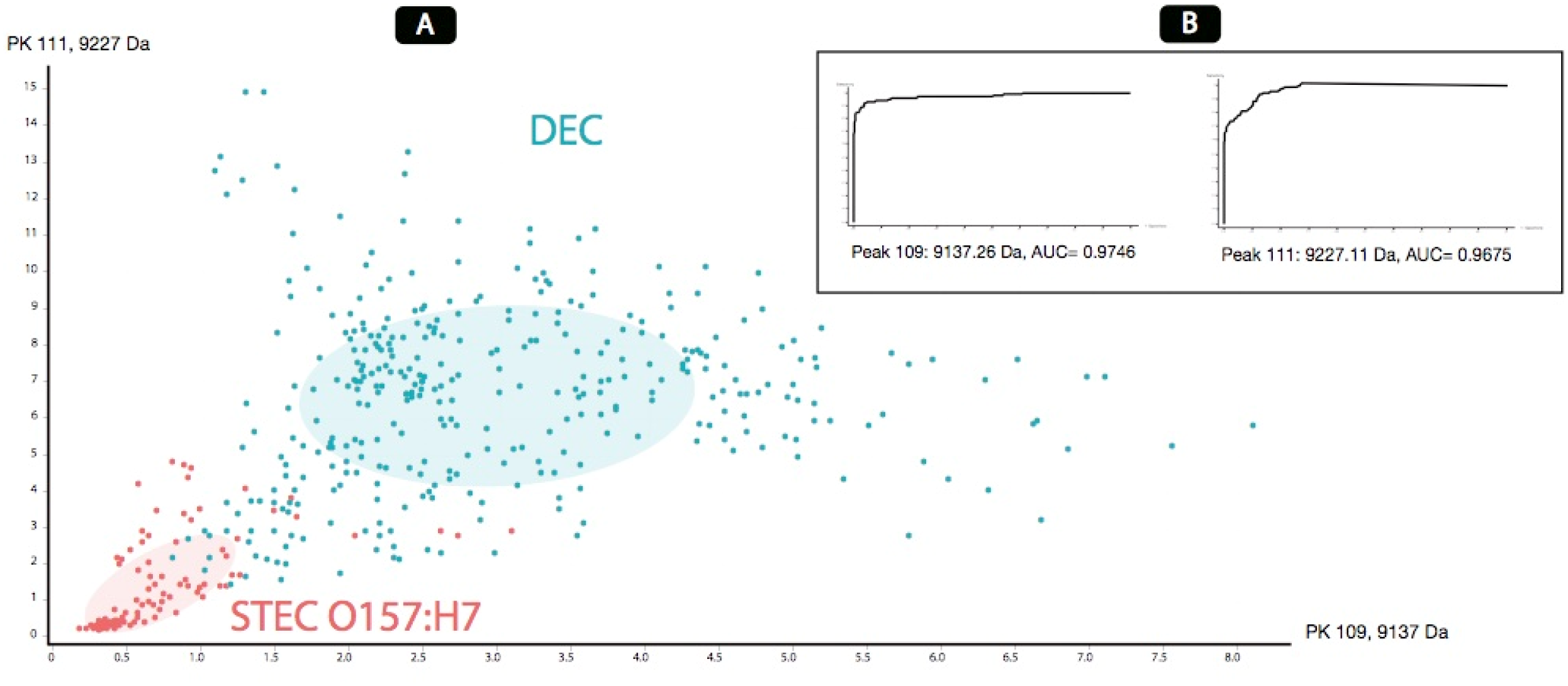
**A-** 2D graph of peak distribution of the 2-class model. **B-** ROC curves of the most discriminating peaks according to the analyses performed.

### Classifier Models

The predictive models were calculated based on three available algorithms: GA/kNN, SNN and QC, the results of the different parameters of each algorithm are summarized in **Table 1**.

**Table 1.** Results of RC, CV and integration areas for each algorithm.

| Classifiers<br>Algorithms | RC | CV | Integration areas used by each<br>model |
| --- | --- | --- | --- |
| **GA / kNN** | 100.00% | 100.00% | 3082;4939;5080;8813;8994 |
| **QC** | 92.83% | 92.47% | 6389;9136;9226 |

RC= Rec ognit ion
Capability; VC=Cross Validation

The SNN algorithm was discarded from the successive analyses due that the results obtained were not optimal.

As a result of the external validation carried out with the set of 142 isolates, a good performance was observed with the GA/kNN algorithm and with the QC algorithm. Nevertheless, we decided to combine both models; first, using the QC algorithm but applying a cut-off value >= 1.55, since from this value 100% correct classifications were observed compared to the reference technique (Figure 3).

**Figure 3.**
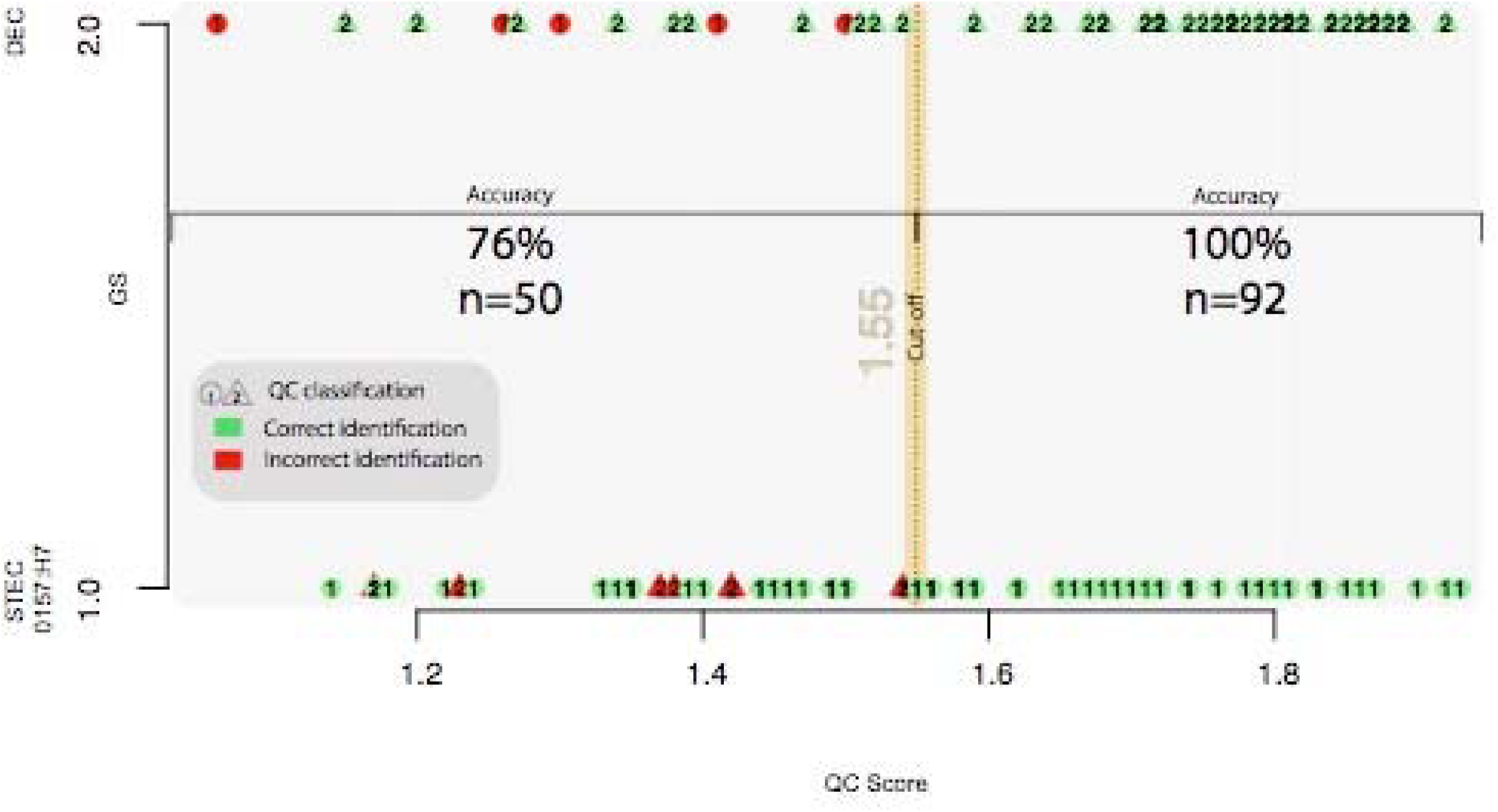
Graph of QC scores where the 100% concordance of the QC algorithm with the reference method is observed from the standardized cut-off value of 1.55.

On the other hand, the isolates that were classified with a QC value <1.55 the GA/kNN algorithm was applied (Figure 4). This combination managed to increase the identification up to 97%, as detailed later in Table 3.

**Figure 4.**
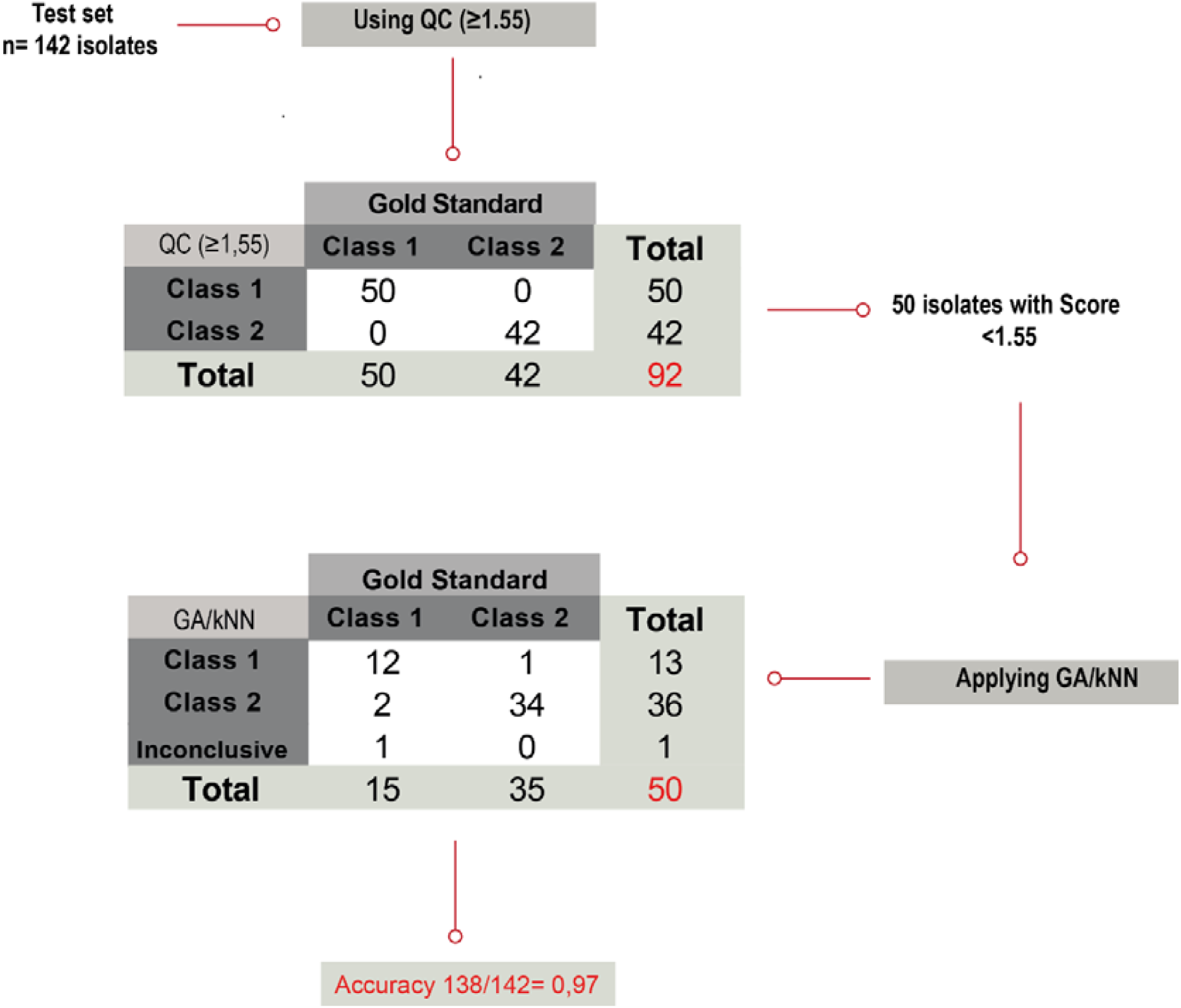
Algorithm applied for the identification of STEC O157:H7 based on predictive classification models.

There were three isolates incorrectly classified using this scheme and one was considered Inconclusive since the result of the QC algorithm was <1.55 and when applying the GA/kNN algorithm dissimilar results were obtained between the sample and the duplicate.

### Biomarker detection

10 differential peaks with statistical significance were found in both software; of which 9 correspond to the STEC O157:H7 pathotype and a single biomarker was present only in NON STEC O157 (Figure 5).

**Figure 5.**
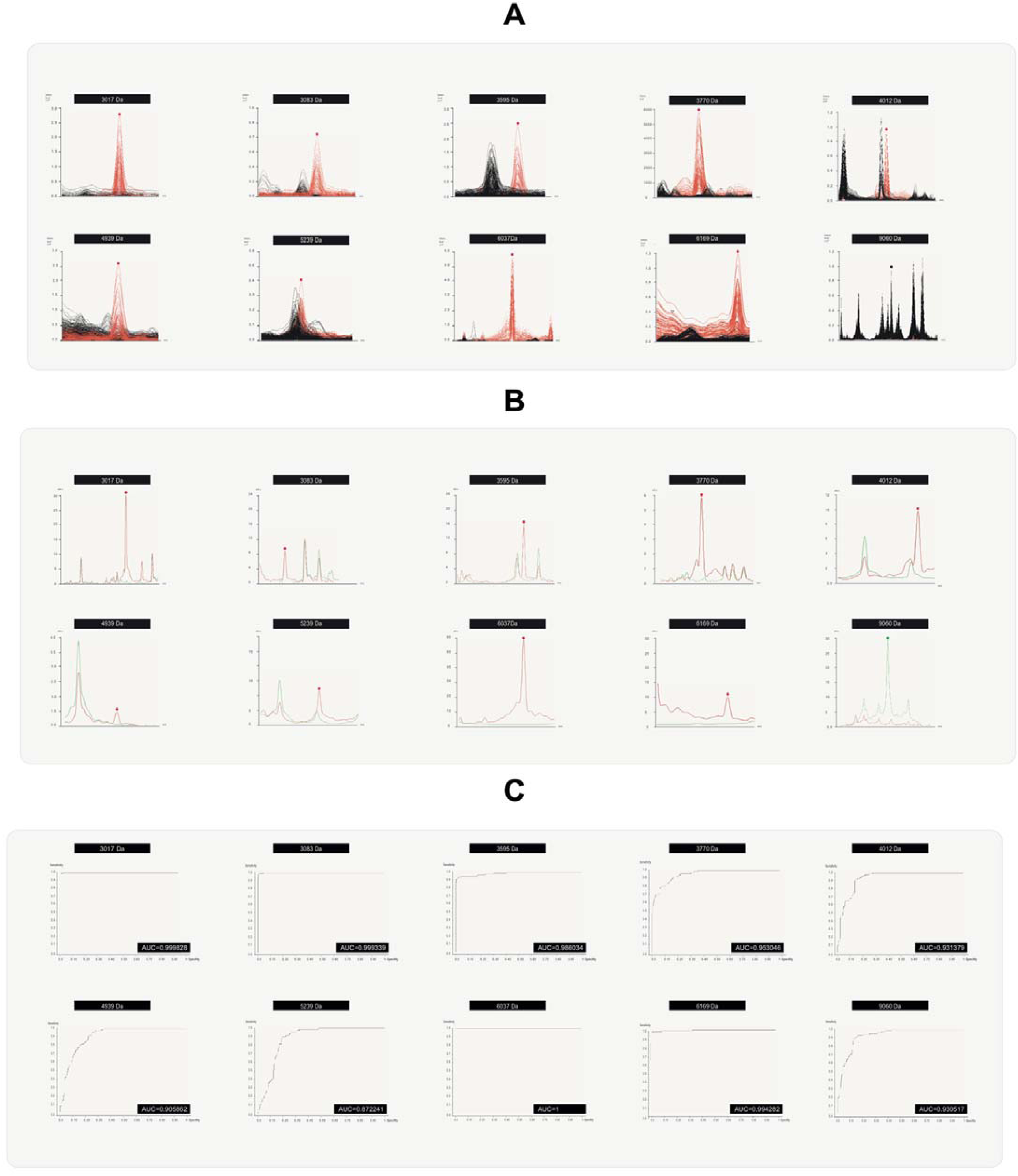
A- Characteristic peaks (biomarkers) in individual spectra of STEC O157:H7 samples (red) versus DEC samples (black), obtained by manual analysis in Flex Analysis v3.4 software. B- Average spectra of the same peaks, STEC O157:H7 (red); DEC (green), obtained by ClinPro Tools v3.0. C- ROC curves and AUC values originated from the external MATLAB software tool integrated in ClinPro Tools.

The profile of the 10 potential biomarkers selected for the differentiation of STEC O157:H7 from DEC isolates is shown in Table 2. The description of the profiles found for all the challenged isolates is found in Table S3 of the Supplementary Material.

**Table 2.** Profile of the 10 potential biomarkers selected for the differentiation of STEC O157:H7 from DEC.

| Classification | Biomarkers (m/z) |  |  |  |  |  |  |  |  |  | 80-100 %<br>70-80 %<br>60-70 %<br><60 %<br>90-100 %<br>80-90 %<br>Presence<br>Absence |
| --- | --- | --- | --- | --- | --- | --- | --- | --- | --- | --- | --- |
|  | 3017 | 3083 | 3595 | 3770 | 4012 | 4939 | 5238 | 6037 | 6169 | 9060 |  |
| **CLASS 1**<br>(O157:H7; n=65) | 72%<br>(n=47) | 45%<br>(n=29) | 51%<br>(n=33) | 85%<br>(n=55) | 40%<br>(n=26) | 38%<br>(n=25) | 97%<br>(n=63) | 86%<br>(n=56) | 55%<br>(n=36) | 100%<br>(n=65) |  |
| **CLASS 2**<br>(DEC; n=77) | 96%<br>(n=74) | 100%<br>(n=77) | 99%<br>(n=76) | 99%<br>(n=76) | 100%<br>(n=77) | 100%<br>(n=77) | 100%<br>(n=77) | 97%<br>(n=75) | 100%<br>(n=77) | 96%<br>(n=74) |  |

Based on the analysis of the results obtained from the BM challenge on 142 new samples in duplicate, it can be confirmed that the absence of the 9060 Da peak, added to the detection of at least one of the other nine peaks described in this work, confirmed the identification of an isolate of STEC O157:H7, although most of these isolates (92%), in addition to not showing the peak at 9060 Da, had 3 or more biomarkers.

One isolate of STEC O157:H7 and 3 other isolates (STEC NON O157, ETEC, and non-toxigenic O157) did not present any of the peaks listed above, and thus could not be classified.

Regarding the NON STEC O157, 96% presented the peak of m/z 9060 Da and 93.5% none of the nine peaks previously described.

Finally, the sensitivity of the search for probable biomarker peaks was 96.9% and the specificity 100%, as shown in Table 3.

**Table 3.** Sensitivity, Specificity, PPV and NPV values of the different approaches evaluated and the statistical relationship between them.

| Evaluated<br>Parameter | Approaches |  |  |  |  |
| --- | --- | --- | --- | --- | --- |
|  | A | B | C | D | E |
|  | Model<br>GA/kNN | Model<br>QC | Combination<br>QC<br>+<br>GA/kNN | Biomarkers | Combination<br>QC + GA/kNN<br>+<br>Biomarkers |
| **Sensitivity** | 80,00% | 92,30% | 95,40% | 96,90% | 98,50% |
| **Specificity** | 96,10% | 90,90% | 98,70% | 100% | 100% |
| **PPV** | 94,50% | 89,60% | 98,40% | 100% | 100% |
| **NPV** | 85,10% | 93,30% | 96,20% | 97,50% | 98,70% |

**PPV=** Positive predictive value; **NPV=** Negative predictive value

The difference between the sensitivity and specificity of the mathematical model with respect to the manual (C and D) and the two best methods (D and E) and with respect to the reference method was estimated to conclude whether these differences were statistically significant or not. In this way, if the 95% confidence interval of the differences contains the value 0, it is concluded that there are no statistically significant differences, otherwise there are differences. The results of this analysis are detailed in Table 4.

**Table 4.** Results of the difference between the sensitivity and specificity of the mathematical and manual model and of the two best methods with respect to the reference method.

| Combination<br>of approaches | Difference<br>sensitivity (%) | Difference<br>specificity (%) | Confidence<br>Intervals<br>95% |  | Conclusion |
| --- | --- | --- | --- | --- | --- |
|  |  |  | Lower | Upper |  |
| **C vs D** | -1.54 | - | -8.58 | 6.1 | The difference between the sensitivity of the methods **is not statistically significant** |
|  | - | 1.3 | -3.57 | 7.00 | The difference between the sensitivity of the methods **is not statistically significant** |
| **D vs E** | 1.56 | - | -4.52 | 9.03 | The difference between the sensitivity of the methods **is not statistically significant** |
|  | - | 0 | -5.63 | 4.87 | The difference between the sensitivity of the methods **is not statistically significant** |

Therefore, despite the fact that there were no statistically significant differences between the performance values of approaches D and E, the benefits of their combined use are evident, as described in numerous publications in the current literature **[15]** and it follows from this work in the resolution of discordant cases.

In summary, if both developments are applied in a complementary way to isolates that could not be correctly classified by automated training or did not present any of the peaks considered biomarkers, accurate detection of 98.5% of STEC O157:H7 isolates is achieved and the correct classification of 99.3% of all the isolates studied

It was observed that 3/3 isolates incorrectly classified by the predictive models were correctly resolved by the BM finding method and 3/4 that could not be classified because they did not present the peaks, were resolved by mathematical models; a single case could not be resolved by either of the two methods, reaffirming the usefulness of the combined use of both approaches (Table 5).

**Table 5.** Discordant cases of the different approaches compared with the gold standard results.

| Samples ID | GA/kNN+QC | Biomarkers | GA/kNN + QC<br>+<br>Biomarkers | Gold<br>standard |
| --- | --- | --- | --- | --- |
| 750/18 | Class 2 | Class 1 | Class 1 | Class 1 |
| 506/18 | Class 2 | N/D | Class 2 | Class 2 |
| 385/16 | Class 1 | Class 2 | Class 2 | Class 2 |
| 504/18 | Class 2 | Class 1 | Class 1 | Class 1 |
| 329/18 | Class 2 | N/D | Class 2 | Class 2 |
| 714/18 | N/D | N/D | N/D | Class 1 |
| 493/18 | Class 2 | N/D | Class 2 | Class 2 |

N/D=not determinated

The table of the results obtained on the total number of isolates applying machine learning and biomarker detection, in comparison with current reference techniques, can be found in Table S3 the Supplementary Material.

## CONCLUSIONS

Due to the important analytical capabilities that MS currently has, added to the speed of results and lower-cost, the possible implementation of the MALDI-TOF MS system coupled to simple and practical artificial intelligence tools could be considered as a STEC O157:H7 diagnostic screening method.

Through the proteomic analysis of the information contained in the spectra of the different classes of *E. coli*, and applying a combination of predictive models based on machine learning, it was possible to quickly identify 94% of the STEC O157:H7 isolates and precise, starting from characteristic suspicious colonies, which implied a substantial saving of time and resources in the routine of the conventional laboratory. By combining this approach with the search for potential biomarker peaks, the percentage of correct identifications rose to 98.5%.

There were several previous attempts in the literature to detect STEC O157 by MALDI-TOF Mass Spectrometry **[16-19]**, however, no defining peaks were found in any of the previous works. and without the requirement of complex extraction techniques or equipment with greater discriminatory power, such as the TOF-TOF type, or peak readings above 10,000 Da, which are generally less detected. On the other hand, here we detect a large number of reproducible peaks in the reading range of the order of 3000 to 9000Da by direct method, without the need to make any modifications, which results in a simple, fast and easily transferable procedure to less complex clinical laboratories that have the technology.

In some cases, a difference in the presence of a peak was observed in the duplicate of the same sample, which may be due to operator errors or by using the direct method, which presents greater variability than the extraction techniques. Evidence from the literature suggests that the protein extraction method extends or improves the range of peaks identified **[16,20]**. However, in this work direct method was prioritized, because it is much simpler, faster and easily applicable in the routine of any clinical laboratory. Due to the variability of the method evidenced on some challenged spectra, the importance of working with technical replicates and analysing the presence/absence of several characteristic peaks is highlighted, in order to reduce the risk of making errors during the classification of one class or another. It is also evident that the search for a single peak is not enough, but rather the joint analysis of a profile, either manually or automatically, in order to performed a reliable identification.

According to previous works by Mazzeo, 2006 and Teruyo, 2014, it was possible to confirm that the absence of the peak at 9060 Da is a useful indicator of STEC O157:H7, although it is not definitive per se, as could be evidenced in our work, where the finding of one or more of the nine detected peaks would confirm the presence of a STEC O157:H7 isolate, because these are not normally found in the other diarrheal types of the genus.

The total absence of biomarker peaks that occurred in 4 isolates (three of them NON STEC O157 that would generally present a single peak) may also be due to errors in the technical procedure, which, in these cases, would be convenient to repeat or confirm by other methods if only this approach is available.

A particular case occurred where the presence of 4 peaks was detected, as occurs on isolate of STEC O157, but with the presence, in addition to the representative peak of the DEC group at m/z 9060 Da, this isolate corresponded to a O157 H7 non-toxigenic type, which had been isolated from a meat processing plant, this strengthens the idea of being in the presence of a STEC isolate O157:H7 with the loss of the Shiga toxin phage, in the same way as described in other works **[21-22]**. Said isolate also presented other virulence factors such as enterohemolysin and eae, typical of STEC O157:H7.

Finally, the approach based on the detection of the presence/absence of peaks, although it is a manual method that requires a longer analysis time, presented excellent performance values and the were no differences regarding in sensitivity and specificity compared to the mathematical model, added to the availability of the Flex Analysis software in the equipment, the detection of these biomarker peaks could be applied in laboratories as a rapid screening method for suspicious STEC O157:H7 isolates.

## REFERENCES

1. Rivas, M., Miliwebsky, E., Chinen, I., Deza, N., & Leotta, G. A. (2006). Epidemiología del síndrome urémico hemolítico en Argentina. Diagnóstico del agente etiológico, reservorios y vías de transmisión. Medicina, 66(supplement 3), 27–32.

2. Miliwebsky E, Schelotto F, Varela G, Luz D, Chinen I, Piazza RMF. Human Diarrheal Infections: Diagnosis of Diarrheagenic Escherichia coli Pathotypes Escherichia coli in the Americas. Torres AG (ed.). En: Escherichia coli in the Americas. Springer E-Book. ISBN: 978-3-319-45092-6. 2016. Chapter 15. p. 343–69.

3. Kallow, W., Erhard, M., Shah, H.N., Raptakis, E. and Welker, M. (2010). MALDI-TOF MS for Microbial Identification: Years of Experimental Development to an Established Protocol. In Mass Spectrometry for Microbial Proteomics (eds H.N. Shah and S.E. Gharbia). https://doi.org/10.1002/9780470665497.ch12

4. He Y, Li H, Lu X, Stratton CW, Tang YW. Mass spectrometry biotyper system identifies enteric bacterial pathogens directly from colonies grown on selective stool culture media. J Clin Microbiol. 2010;48(11):3888–3892. doi:10.1128/JCM.01290-10

5. Cherkaoui A, Hibbs J, Emonet S, et al. Comparison of two matrix-assisted laser desorption ionization-time of flight mass spectrometry methods with conventional phenotypic identification for routine identification of bacteria to the species level. J Clin Microbiol. 2010;48(4):1169–1175. doi:10.1128/JCM.01881-09

6. Rocca MF, Zintgraff JC, Dattero ME, et al. A combined approach of MALDI-TOF mass spectrometry and multivariate analysis as a potential tool for the detection of SARS-CoV-2 virus in nasopharyngeal swabs. J Virol Methods. 2020;286:113991. doi:10.1016/j.jviromet.2020.113991

7. Miliwebsky E. Manual de procedimientos Escherichia coli productor de toxina Shiga en el marco de la detección de E. coli diarreigénico. 2019. http://sgc.anlis.gob.ar/handle/123456789/2307

8. Espinal P, Seifert H, Dijkshoorn L, Vila J, Roca I. Rapid and accurate identification of genomic species from the Acinetobacter baumannii (Ab) group by MALDI-TOF MS. Clin Microbiol Infect. 2012;18(11):1097–1103. doi:10.1111/j.1469-0691.2011.03696.x

9. Bruker Daltonik GmbH, 2011. ClinPro Tools User Manual Version 3.0.BrukerDaltonik GmbH, Bremen https://doi.org/10.1371/journal.pone.0230334.

10. Camoez, M., Sierra, J.M., Dominguez, M.A., Ferrer-Navarro, M., Vila, J., Roca, I., 2016. Automated categorization of methicillin-resistant Staphylococcus aureus clinical isolates into different clonal complexes by MALDI-TOF mass spectrometry. Clin.Microbiol. Infect. 22 (2) https://doi.org/10.1016/j.cmi.2015.10.009, 161.e1-161. e7.

11. Zhang, H., Cao, J., Li, L., Liu, Y., Zhao, H., Li, N., Li, B., Zhang, A., Huang, H., Chen, S., Dong, M., Yu, L., Zhang, J., Chen, L., 2015. Identification of urine protein biomarkers with the potential for early detection of lung cancer. Sci.Rep. 5, 11805. https://doi.org/10.1038/srep11805.

12. Wang, H.Y., Lien, F., Liu, T.P., Chen, C.H., Chen, C.J., Lu, J.J., 2018. Application of a MALDI-TOF analysis platform (ClinProTools) for rapid and preliminary report of sequence types in Taiwan. PeerJ. 6, e5784. https://doi.org/10.7717/peerj.5784

13. Stephens, M.A., 1974. EDF statistics for goodness of fit and some comparisons. JASA 69, 730–737.

14. Landis, J., & Koch, G. The measurement of observer agreement for categorical data. Biometrics, 1977; 33, 159–174

15. Yumi Kubo, Osamu Ueda, Sawa Nagamitsu, Hachiro Yamanishi, Akihiro Nakamura, Masaru Komatsu, Novel strategy of rapid typing of Shiga toxin-producing Escherichia coli using MALDI Biotyper and ClinProTools analysis. Journal of Infection and Chemotherapy, Volume 27, Issue 8, 2021, Pages 1137–1142, ISSN 1341-321X, https://doi.org/10.1016/j.jiac.2021.03.002.

16. Mazzeo Maria Fiorella, Alida Sorrentino, Marcello Gaita, Giuseppina Cacace, Michele Di Stasio, Angelo Facchiano, Giuseppe Comi, Antonio Malorni, and Rosa Anna Siciliano Matrix-Assisted Laser Desorption Ionization-Time of Flight Mass Spectrometry for the Discrimination of Food-Borne Microorganisms. Appl Environ Microbiol. 2006 Feb; 72(2): 1180–1189. doi: 10.1128/AEM.72.2.1180-1189.2006.

17. Clark CG, Kruczkiewicz P, Guan C, et al. Evaluation of MALDI-TOF mass spectroscopy methods for determination of Escherichia coli pathotypes. J Microbiol Methods. 2013;94(3):180–191. doi:10.1016/j.mimet.2013.06.020

18. Fagerquist CK, Garbus BR, Miller WG, et al. Rapid identification of protein biomarkers of Escherichia coli O157:H7 by matrix-assisted laser desorption ionization-time-of-flight-time-of-flight mass spectrometry and top-down proteomics. Anal Chem. 2010;82(7):2717–2725. doi:10.1021/ac902455d

19. Ojima-Kato T, Yamamoto N, Suzuki M, Fukunaga T, Tamura H. Discrimination of Escherichia coli O157, O26 and O111 from other serovars by MALDI-TOF MS based on the S10-GERMS method. PLoS One. 2014;9(11):e113458. Published 2014 Nov 20. doi:10.1371/journal.pone.0113458

20. Ochoa ML, Harrington PB. Immunomagnetic isolation of enterohemorrhagic Escherichia coli O157:H7 from ground beef and identification by matrix-assisted laser desorption/ionization time-of-flight mass spectrometry and database searches. Anal Chem. 2005;77(16):5258–5267. doi:10.1021/ac0502596

21. Bielaszewska M, Köck R, Friedrich AW, et al. Shiga toxin-mediated hemolytic uremic syndrome: time to change the diagnostic paradigm?. PLoS One. 2007;2(10):e1024. Published 2007 Oct 10. doi:10.1371/journal.pone.0001024

22. Friedrich AW, Zhang W, Bielaszewska M, et al. Prevalence, virulence profiles, and clinical significance of Shiga toxin-negative variants of enterohemorrhagic Escherichia coli O157 infection in humans. Clin Infect Dis. 2007;45(1):39–45. doi:10.1086/518573

