## Supplementary material for "Rapid and accurate identification of *Escherichia coli* STEC O157:H7 by mass spectrometry, artificial intelligence and detection of specific biomarkers peaks"

**Table S1.** Set of samples used for the development of predictive models and the detection of biomarkers (N=60).

| Sample ID | Type | Shiga Toxin | Sample ID | Type | Shiga Toxin |
| --- | --- | --- | --- | --- | --- |
| 456/18 | **O157:H7** | **2** | 783/12 | EPEC | - |
| 450/18 | O157:H7 | 2 | 1052/10 | EPEC | - |
| 453/18 | O157:H7 | 2 | 545/10 | EPEC | - |
| 379/18 | O157:H7 | 2 | 1145/09 | EPEC | - |
| 404/18 | O157:H7 | 2 | 712/11 | EPEC | - |
| 370/18 | O157:H7 | 2 | 366/11 | EPEC | - |
| 672/17 | O157:H7 | 2 | 972/11 | EPEC | - |
| 340/18 | O157:H7 | 2 | 847/11 | EPEC | - |
| 419/18 | O157:H7 | 2 | 1051/11 | EPEC | - |
| 370/18 | O157:H7 | 2 | 1038/11 | EPEC | - |
| 373/02 | O103H2 | 1/2 | 828/11 | EPEC | - |
| 178/17 | 121H19 | 2 | 712/11 | EPEC | - |
| 370/02 | O113H21 | 2 | 366/11 | EPEC | - |
| 231/09 | O111NM | 1/2 | 330/11 | EPEC | - |
| 433/01 | O111NM | 1/2 | 220/11 | EPEC | - |
| 67/03 | O111NM | 1/2 | 776/12 | ETEC | - |
| 53/01 | O145NM | 2 | 100/13 | ETEC | - |
| 2/02 | O145NM | 2 | 904/02 | ETEC | - |
| 267/01 | O91H21 | 2 | 902/02 | ETEC | - |
| 71/02 | O26H11 | 1 | 901/02 | ETEC | - |
| 208/18 | O26H11 | 1 | 635/05 | ETEC | - |
| 49/18 | O103H2 | 1/2 | 228/05 | ETEC | - |
| 727/17 | 121H19 | 2 | 30/13 | ETEC | - |
| 958/05 | O91H21 | 2 | 909/02 | ETEC | - |
| 709/10 | O113 H2 | 2 | 946/02 | ETEC | - |
| 582/10 | O174H21 | 2 | 985/02 | ETEC | - |
| 357/18 | O145NM | 2 | 696/10 | ETEC | - |
| 653/17 | O26H11 | 1 | D18 | ETEC | - |
| T154 | O103H2 | 1/2 | 07/03 | ETEC | - |
| T171 | O91H21 | 2 | 823/11 | ETEC | - |

**Table S2.** List of the best peaks found in Clin ProTools based on the statistical parameters

| **Index** | **Mass** | **DAve** | **PTTA** | **PWKW** | **PAD** |
| --- | --- | --- | --- | --- | --- |
| 109 | 9137.26 | 2.52 | < 0.000001 | < 0.000001 | < 0.000001 |
| 111 | 9227.11 | 5.68 | < 0.000001 | < 0.000001 | < 0.000001 |
| 82 | 6388.77 | 2.86 | < 0.000001 | < 0.000001 | < 0.000001 |
| 110 | 9192.2 | 2.86 | < 0.000001 | < 0.000001 | < 0.000001 |
| 84 | 6487.52 | 1.87 | < 0.000001 | < 0.000001 | 0.000107 |
| 112 | 9270.87 | 1.36 | < 0.000001 | < 0.000001 | < 0.000001 |
| 50 | 4614.88 | 6.83 | < 0.000001 | < 0.000001 | < 0.000001 |
| 118 | 10139.3 | 2.52 | < 0.000001 | < 0.000001 | 0.000829 |
| 108 | 9064.73 | 20.22 | < 0.000001 | < 0.000001 | < 0.000001 |
| 49 | 4533.61 | 11.7 | < 0.000001 | < 0.000001 | < 0.000001 |
| 46 | 4439.65 | 6.38 | < 0.000001 | < 0.000001 | 0.000806 |
| 121 | 10470.16 | 6.81 | < 0.000001 | 0 | < 0.000001 |
| 113 | 9537.37 | 11.18 | < 0.000001 | < 0.000001 | < 0.000001 |
| 106 | 8876.88 | 4.35 | < 0.000001 | < 0.000001 | < 0.000001 |
| 107 | 8994.92 | 3.86 | < 0.000001 | < 0.000001 | < 0.000001 |
| 83 | 6412.45 | 4.52 | < 0.000001 | < 0.000001 | 0.000697 |
| 51 | 4769.98 | 10.71 | < 0.000001 | < 0.000001 | < 0.000001 |
| 124 | 10693.69 | 1.15 | < 0.000001 | < 0.000001 | < 0.000001 |
| 85 | 6507.24 | 1.76 | < 0.000001 | < 0.000001 | < 0.000001 |
| 105 | 8815.25 | 1.42 | < 0.000001 | 0 | < 0.000001 |
| 27 | 3184.46 | 2.11 | < 0.000001 | < 0.000001 | 0.00000603 |
| 47 | 4449.77 | 3.49 | < 0.000001 | < 0.000001 | 0.00195 |
| 134 | 12969.7 | 0.63 | < 0.000001 | < 0.000001 | < 0.000001 |
| 22 | 3083.47 | 8.67 | < 0.000001 | 0 | < 0.000001 |
| 91 | 7233.86 | 1.73 | < 0.000001 | < 0.000001 | < 0.000001 |
| 71 | 6038.47 | 89.71 | < 0.000001 | 0 | < 0.000001 |
| 52 | 4778.66 | 9.9 | < 0.000001 | < 0.000001 | < 0.000001 |
| 28 | 3205.27 | 2.23 | < 0.000001 | < 0.000001 | < 0.000001 |
| 133 | 12769.38 | 0.77 | < 0.000001 | < 0.000001 | < 0.000001 |
| 119 | 10299.85 | 3.19 | < 0.000001 | < 0.000001 | 0.00167 |
| 114 | 9611.74 | 2.28 | < 0.000001 | < 0.000001 | < 0.000001 |
| 126 | 11203.18 | 1.33 | < 0.000001 | < 0.000001 | < 0.000001 |
| 122 | 10538.31 | 1.96 | < 0.000001 | 0 | < 0.000001 |
| 21 | 3017.83 | 36.78 | < 0.000001 | 0 | < 0.000001 |
| 48 | 4498.63 | 4.54 | < 0.000001 | < 0.000001 | 0.000142 |
| 1 | 2008.92 | 1.95 | < 0.000001 | 0 | < 0.000001 |
| 59 | 5238.79 | 5.15 | < 0.000001 | 0 | < 0.000001 |
| 66 | 5602.91 | 1.77 | < 0.000001 | < 0.000001 | < 0.000001 |
| 75 | 6169.63 | 19.73 | < 0.000001 | 0 | < 0.000001 |
| 72 | 6079.8 | 11.58 | < 0.000001 | 0 | < 0.000001 |
| 43 | 4365.28 | 12.72 | < 0.000001 | < 0.000001 | < 0.000001 |
| 42 | 4349.49 | 1.71 | < 0.000001 | < 0.000001 | < 0.000001 |
| 55 | 5082.5 | 3.67 | < 0.000001 | 0 | < 0.000001 |
| 20 | 3003.44 | 2.15 | < 0.000001 | 0 | < 0.000001 |
| 127 | 11220.65 | 1.39 | < 0.000001 | < 0.000001 | < 0.000001 |
| 81 | 6358.84 | 1.46 | < 0.000001 | < 0.000001 | < 0.000001 |
| 36 | 3936.46 | 3.16 | < 0.000001 | < 0.000001 | < 0.000001 |
| 5 | 2179.44 | 2.89 | < 0.000001 | < 0.000001 | < 0.000001 |
| 74 | 6153.6 | 5.76 | < 0.000001 | 0 | < 0.000001 |
| 58 | 5152.09 | 5.12 | < 0.000001 | < 0.000001 | < 0.000001 |
| 44 | 4379.9 | 3.55 | < 0.000001 | < 0.000001 | < 0.000001 |
| 73 | 6116.73 | 4.68 | < 0.000001 | 0 | < 0.000001 |
| 139 | 15315.11 | 0.46 | < 0.000001 | < 0.000001 | < 0.000001 |
| 70 | 5936.55 | 6.36 | < 0.000001 | 0 | < 0.000001 |
| 132 | 12653.54 | 0.39 | < 0.000001 | < 0.000001 | < 0.000001 |
| 56 | 5097.6 | 22.85 | < 0.000001 | 0 | < 0.000001 |
| 67 | 5613.95 | 2.16 | < 0.000001 | < 0.000001 | < 0.000001 |
| 99 | 7871.91 | 3.43 | < 0.000001 | < 0.000001 | 0.189 |
| 41 | 4211.68 | 3.83 | < 0.000001 | 0 | < 0.000001 |
| 125 | 10745.29 | 0.5 | < 0.000001 | < 0.000001 | < 0.000001 |
| 26 | 3178.45 | 1.25 | < 0.000001 | < 0.000001 | < 0.000001 |
| 68 | 5727.82 | 2.03 | < 0.000001 | < 0.000001 | 0.0000581 |
| 120 | 10388.14 | 0.67 | < 0.000001 | 0 | < 0.000001 |
| 93 | 7334.35 | 1.67 | < 0.000001 | < 0.000001 | < 0.000001 |
| 17 | 2965.76 | 1.52 | < 0.000001 | 0 | < 0.000001 |
| 69 | 5873.32 | 4.96 | < 0.000001 | 0 | < 0.000001 |
| 92 | 7275.01 | 9.98 | < 0.000001 | < 0.000001 | 0.0000279 |
| 19 | 2997.53 | 1.09 | < 0.000001 | 0 | < 0.000001 |
| 97 | 7661.83 | 0.82 | < 0.000001 | < 0.000001 | < 0.000001 |
| 128 | 11449.21 | 0.83 | < 0.000001 | < 0.000001 | < 0.000001 |
| 89 | 7158.06 | 6.31 | < 0.000001 | < 0.000001 | < 0.000001 |
| 33 | 3596.14 | 17.55 | < 0.000001 | 0 | < 0.000001 |
| 31 | 3443.6 | 2.52 | < 0.000001 | 0 | < 0.000001 |
| 15 | 2935.49 | 1.39 | < 0.000001 | 0 | < 0.000001 |
| 62 | 5382.18 | 22.44 | < 0.000001 | 0 | < 0.000001 |
| 76 | 6213.28 | 2.7 | < 0.000001 | 0 | < 0.000001 |
| 34 | 3637.46 | 5.59 | < 0.000001 | < 0.000001 | 0.000979 |
| 10 | 2546.29 | 5.01 | < 0.000001 | 0 | < 0.000001 |
| 140 | 15405.58 | 0.42 | < 0.000001 | < 0.000001 | < 0.000001 |
| 29 | 3423.52 | 3.3 | < 0.000001 | 0 | < 0.000001 |
| 38 | 4012.83 | 9.93 | < 0.000001 | 0 | < 0.000001 |
| 35 | 3771.26 | 7.84 | < 0.000001 | 0 | < 0.000001 |
| 90 | 7200.97 | 0.87 | < 0.000001 | < 0.000001 | < 0.000001 |
| 95 | 7421.3 | 0.71 | < 0.000001 | < 0.000001 | < 0.000001 |
| 54 | 5072.79 | 2.05 | < 0.000001 | < 0.000001 | < 0.000001 |
| 11 | 2688.9 | 6.84 | < 0.000001 | < 0.000001 | < 0.000001 |
| 101 | 8007.36 | 3.56 | < 0.000001 | < 0.000001 | 0 |
| 141 | 15723.48 | 0.21 | < 0.000001 | < 0.000001 | < 0.000001 |
| 144 | 17464.78 | 0.13 | < 0.000001 | < 0.000001 | < 0.000001 |
| 60 | 5340.7 | 1.78 | < 0.000001 | < 0.000001 | < 0.000001 |
| 143 | 16009.85 | 0.15 | < 0.000001 | < 0.000001 | < 0.000001 |
| 135 | 13650.09 | 0.15 | < 0.000001 | < 0.000001 | < 0.000001 |
| 87 | 6826.88 | 0.46 | < 0.000001 | < 0.000001 | 0.000025 |
| 100 | 7928.45 | 0.48 | < 0.000001 | < 0.000001 | < 0.000001 |
| 86 | 6685.34 | 0.42 | < 0.000001 | 0.0133 | < 0.000001 |
| 145 | 18153.95 | 0.52 | < 0.000001 | < 0.000001 | < 0.000001 |
| 57 | 5114.88 | 0.85 | < 0.000001 | < 0.000001 | < 0.000001 |
| 12 | 2700.85 | 0.44 | < 0.000001 | < 0.000001 | < 0.000001 |
| 64 | 5420.57 | 0.89 | < 0.000001 | < 0.000001 | < 0.000001 |
| 24 | 3149.42 | 0.53 | < 0.000001 | 0.00709 | < 0.000001 |
| 131 | 12215.53 | 0.23 | < 0.000001 | 0.113 | < 0.000001 |
| 137 | 14705.01 | 0.15 | < 0.000001 | < 0.000001 | < 0.000001 |
| 6 | 2185.24 | 1.13 | 0.00000147 | < 0.000001 | < 0.000001 |
| 45 | 4408.22 | 0.63 | 0.00000253 | < 0.000001 | < 0.000001 |
| 16 | 2958.07 | 0.43 | 0.00000676 | 0.00133 | < 0.000001 |
| 123 | 10652.72 | 0.24 | 0.0000117 | 0.00000217 | 0.00000177 |
| 25 | 3157.44 | 1.46 | 0.0000143 | 0.000315 | 0.000497 |
| 103 | 8370.14 | 2.02 | 0.000034 | 0.0000302 | < 0.000001 |
| 53 | 4871.85 | 6.65 | 0.000125 | 0.000563 | < 0.000001 |
| 18 | 2980.33 | 1.69 | 0.000154 | 0.000257 | < 0.000001 |
| 63 | 5399.37 | 0.92 | 0.000181 | 0.0000148 | < 0.000001 |
| 96 | 7466.02 | 0.21 | 0.000577 | 0.0311 | < 0.000001 |
| 136 | 14361.14 | 0.05 | 0.000989 | 0.0258 | < 0.000001 |
| 3 | 2081.96 | 0.19 | 0.00133 | 0.036 | < 0.000001 |
| 142 | 15843.63 | 0.08 | 0.00226 | 0.0123 | < 0.000001 |
| 98 | 7709.28 | 0.96 | 0.00226 | 0.00335 | < 0.000001 |
| 79 | 6301.7 | 1.2 | 0.00229 | 0.526 | < 0.000001 |
| 115 | 9739.97 | 6.81 | 0.00415 | 0.00143 | < 0.000001 |
| 77 | 6256.51 | 4.38 | 0.00439 | 0.000563 | < 0.000001 |
| 40 | 4185.86 | 1.48 | 0.00537 | 0.0000875 | 0.0918 |
| 13 | 2788.94 | 0.26 | 0.00758 | 0.0000525 | 0.00193 |
| 32 | 3578.85 | 0.91 | 0.0265 | 0.00225 | < 0.000001 |
| 7 | 2413.29 | 0.36 | 0.0374 | 0.317 | < 0.000001 |
| 23 | 3127.07 | 0.85 | 0.0967 | 0.0973 | 0.0000221 |
| 80 | 6317.23 | 1.14 | 0.123 | 0.601 | < 0.000001 |
| 30 | 3434.66 | 0.28 | 0.13 | 0.314 | < 0.000001 |
| 78 | 6272.55 | 0.56 | 0.135 | 0.555 | < 0.000001 |
| 116 | 9809.41 | 0.44 | 0.157 | 0.195 | < 0.000001 |
| 61 | 5350.95 | 0.36 | 0.172 | 0.053 | < 0.000001 |
| 2 | 2056.58 | 0.23 | 0.232 | 0.00394 | < 0.000001 |
| 65 | 5440.23 | 0.16 | 0.308 | 0.013 | < 0.000001 |
| 104 | 8445.89 | 0.09 | 0.388 | 0.384 | 0.0000768 |
| 138 | 14853.05 | 0.01 | 0.511 | 0.902 | < 0.000001 |
| 130 | 12179.74 | 0.03 | 0.564 | 0.0000851 | < 0.000001 |
| 4 | 2102.33 | 0.05 | 0.615 | 0.889 | < 0.000001 |
| 8 | 2429.18 | 0.13 | 0.672 | 0.0198 | < 0.000001 |
| 14 | 2833.09 | 0.28 | 0.718 | 0.264 | < 0.000001 |
| 102 | 8327.33 | 0.23 | 0.797 | 0.00244 | < 0.000001 |
| 39 | 4164.31 | 0.21 | 0.797 | 0.157 | < 0.000001 |
| 129 | 11783.92 | 0.01 | 0.848 | 0.331 | < 0.000001 |
| 117 | 9882.16 | 0.03 | 0.859 | 0.497 | < 0.000001 |
| 9 | 2453.15 | 0.03 | 0.862 | 0.0392 | < 0.000001 |
| 94 | 7396.69 | 0.01 | 0.862 | 0.321 | < 0.000001 |
| 88 | 6857.87 | 0.02 | 0.877 | 0.338 | < 0.000001 |
| 146 | 19474.9 | 0 | 0.895 | 0.221 | < 0.000001 |
| 37 | 4003.52 | 0.02 | 0.973 | 0 | 0 |

DAve=Difference between the maximum and minimum intensity of the average peak of all classes;

PTTA= p-value obtained through the t-test, range 0-1; where 0: good and 1: bad

VPWKW= p value obtained using the Wilcoxon / Kruskal-Wallis test; range 0-1; where 0: good and 1: bad

PAD= p-value obtained by the Anderson-Darling test: range 0-1; 0: non-normal distribution, 1: normal distribution.

**Table S3:** Profiles of the 142 isolates with the binarized values of the different approaches and the presence or absence of the 10 peaks analysed. 1 corresponds to STEC O157, 0 to DEC or not detected. PEAKS: 1 corresponds to the presence of the corresponding peak, 0 absence**.**

| **Sample ID** | **Type** | **Shiga**  **Toxin** | **Source** | **Gold**  **Standard** | **QC** | **GA/kNN** | **QC+GA/kNN** | **BM** | **QC+GA/kNN**  **+**  **BM** | **Biomarkers** | | | | | | | | | |
| --- | --- | --- | --- | --- | --- | --- | --- | --- | --- | --- | --- | --- | --- | --- | --- | --- | --- | --- | --- |
|  |  |  |  |  |  |  |  |  |  | **9060** | **3017** | **3083** | **3595** | **3770** | **4012** | **6037** | **6169** | **4939** | **5238°** |
| 763/18 | O157:H7 | 2 | Clinical | 1 | 1 | 1 | 1 | 1 | 1 | 0 | 1 | 1 | 1 | 0 | 0 | 1 | 1 | 0 | 1 |
| 661/18 | O157:NM | NT | Clinical | 0 | 0 | 0 | 0 | 0 | 0 | 1 | 0 | 0 | 0 | 0 | 0 | 0 | 0 | 0 | 0 |
| 577/18 col 2 | O112:H2 | 2 | Clinical | 0 | 0 | 0 | 0 | 0 | 0 | 1 | 0 | 0 | 0 | 0 | 0 | 0 | 0 | 0 | 0 |
| 569/18 | O145:NM | 2 | Clinical | 0 | 0 | 0 | 0 | 0 | 0 | 1 | 0 | 0 | 0 | 0 | 0 | 0 | 0 | 0 | 0 |
| 464/18 | O157:H16 | NT | Food | 0 | 0 | 0 | 0 | 0 | 0 | 1 | 0 | 0 | 0 | 0 | 0 | 0 | 0 | 0 | 0 |
| 294/18 | O157:H16 | NT | Food | 0 | 0 | 0 | 0 | 0 | 0 | 1 | 0 | 0 | 0 | 0 | 0 | 0 | 0 | 0 | 0 |
| 205/18 | O157:H21 | NT | Food | 0 | 0 | 0 | 0 | 0 | 0 | 1 | 0 | 0 | 0 | 0 | 0 | 0 | 0 | 0 | 0 |
| 135/18 | O157:H16 | NT | Food | 0 | 0 | 0 | 0 | 0 | 0 | 1 | 0 | 0 | 0 | 0 | 0 | 0 | 0 | 0 | 0 |
| 130/18 | O157:H29 | NT | Food | 0 | 0 | 0 | 0 | 0 | 0 | 1 | 0 | 0 | 0 | 0 | 0 | 0 | 0 | 0 | 0 |
| 750/18 | O157:H7 | 2 | Clinical | 1 | 1 | 0 | 0 | 1 | 1 | 0 | 0 | 0 | 1 | 0 | 0 | 0 | 0 | 0 | 0 |
| 751/18 | O157:H7 | 2 | Clinical | 1 | 1 | 1 | 1 | 1 | 1 | 0 | 1 | 1 | 0 | 0 | 0 | 1 | 0 | 0 | 1 |
| 752/18 | O157:H7 | 2 | Clinical | 1 | 1 | 1 | 1 | 1 | 1 | 0 | 1 | 1 | 1 | 0 | 0 | 1 | 1 | 0 | 1 |
| 753/18 | O157:H7 | 2 | Clinical | 1 | 1 | 1 | 1 | 1 | 1 | 0 | 1 | 1 | 1 | 0 | 0 | 1 | 1 | 0 | 1 |
| 762/18 | O157:H7 | 2 | Clinical | 1 | 1 | 1 | 1 | 1 | 1 | 0 | 1 | 1 | 1 | 0 | 0 | 1 | 1 | 0 | 1 |
| 606/18 | O157:H7 | 2 | Clinical | 1 | 1 | 1 | 1 | 1 | 1 | 0 | 1 | 0 | 1 | 1 | 1 | 1 | 0 | 0 | 1 |
| 605/18 | O157:H7 | 2 | Clinical | 1 | 1 | 1 | 1 | 1 | 1 | 0 | 1 | 0 | 1 | 1 | 1 | 1 | 0 | 0 | 1 |
| 567/18 | O157:H7 | 2 | Clinical | 1 | 1 | 1 | 1 | 1 | 1 | 0 | 1 | 0 | 1 | 1 | 1 | 1 | 0 | 0 | 1 |
| 579/18 | O157:H7 | 2 | Clinical | 1 | 1 | 1 | 1 | 1 | 1 | 0 | 1 | 0 | 1 | 1 | 1 | 1 | 1 | 0 | 1 |
| 636/18 | O157:H7 | 2 | Clinical | 1 | 1 | 1 | 1 | 1 | 1 | 0 | 1 | 0 | 1 | 1 | 1 | 1 | 1 | 0 | 1 |
| 662/18 | O157:H7 | 2 | Clinical | 1 | 1 | 1 | 1 | 1 | 1 | 0 | 1 | 0 | 1 | 1 | 1 | 1 | 1 | 0 | 1 |
| 577/18 col1 | O157:H7 | 2 | Clinical | 1 | 1 | 1 | 1 | 1 | 1 | 0 | 1 | 0 | 1 | 0 | 0 | 1 | 0 | 1 | 1 |
| 596/18 | O157:H7 | 2 | Clinical | 1 | 1 | 1 | 1 | 1 | 1 | 0 | 1 | 0 | 1 | 0 | 0 | 1 | 0 | 1 | 1 |
| 573/18 | O145:NM | 2 | Clinical | 0 | 0 | 0 | 0 | 0 | 0 | 1 | 0 | 0 | 0 | 0 | 0 | 0 | 0 | 0 | 0 |
| 494/18 | NT:H19 | 2 | Food | 0 | 0 | 0 | 0 | 0 | 0 | 1 | 1 | 0 | 0 | 0 | 0 | 1 | 0 | 0 | 0 |
| 560/18 | O121:H19 | 2 | Clinical | 0 | 0 | 0 | 0 | 0 | 0 | 1 | 0 | 0 | 0 | 0 | 0 | 0 | 0 | 0 | 0 |
| 554/18 | O145:NM | 2 | Clinical | 0 | 1 | 0 | 0 | 0 | 0 | 1 | 0 | 0 | 0 | 0 | 0 | 0 | 0 | 0 | 0 |
| 527/18 | O145:NM | 2 | Clinical | 0 | 0 | 0 | 0 | 0 | 0 | 1 | 0 | 0 | 0 | 0 | 0 | 0 | 0 | 0 | 0 |
| 506/18 | O145:NM | 2 | Clinical | 0 | 0 | 0 | 0 | 0 | 0 | 0 | 0 | 0 | 0 | 0 | 0 | 0 | 0 | 0 | 0 |
| 93/17 | O103:H3 | 1 | Clinical | 0 | 0 | 0 | 0 | 0 | 0 | 1 | 0 | 0 | 0 | 0 | 0 | 0 | 0 | 0 | 0 |
| 658/18 | O111:NM | 1 | Clinical | 0 | 1 | 0 | 0 | 0 | 0 | 1 | 0 | 0 | 0 | 0 | 0 | 0 | 0 | 0 | 0 |
| 767/17 | ETEC | NT | Clinical | 0 | 0 | 0 | 0 | 0 | 0 | 1 | 0 | 0 | 0 | 0 | 0 | 0 | 0 | 0 | 0 |
| 331/18 | ETEC | NT | Clinical | 0 | 0 | 0 | 0 | 0 | 0 | 1 | 0 | 0 | 0 | 0 | 0 | 0 | 0 | 0 | 0 |
| 427/17 | ETEC | NT | Clinical | 0 | 0 | 0 | 0 | 0 | 0 | 1 | 0 | 0 | 0 | 0 | 0 | 0 | 0 | 0 | 0 |
| 395/18 | ETEC | NT | Clinical | 0 | 0 | 0 | 0 | 0 | 0 | 1 | 0 | 0 | 0 | 0 | 0 | 0 | 0 | 0 | 0 |
| 315/18 | ETEC | NT | Clinical | 0 | 0 | 0 | 0 | 0 | 0 | 1 | 0 | 0 | 0 | 0 | 0 | 0 | 0 | 0 | 0 |
| 230/18 | ETEC | NT | Clinical | 0 | 0 | 0 | 0 | 0 | 0 | 1 | 0 | 0 | 0 | 0 | 0 | 0 | 0 | 0 | 0 |
| 128/18 | EPEC | NT | Clinical | 0 | 0 | 0 | 0 | 0 | 0 | 1 | 0 | 0 | 0 | 0 | 0 | 0 | 0 | 0 | 0 |
| 518/17 | EPEC | NT | Clinical | 0 | 0 | 0 | 0 | 0 | 0 | 1 | 0 | 0 | 0 | 0 | 0 | 0 | 0 | 0 | 0 |
| 823/11 | EPEC | NT | Clinical | 0 | 0 | 0 | 0 | 0 | 0 | 1 | 0 | 0 | 0 | 0 | 0 | 0 | 0 | 0 | 0 |
| 125/18 | ETEC | NT | Clinical | 0 | 0 | 0 | 0 | 0 | 0 | 1 | 0 | 0 | 0 | 1 | 0 | 0 | 0 | 0 | 0 |
| 127/18 | ETEC | NT | Clinical | 0 | 0 | 0 | 0 | 0 | 0 | 1 | 1 | 0 | 0 | 0 | 0 | 0 | 0 | 0 | 0 |
| 746/17 | ETEC | NT | Clinical | 0 | 0 | 0 | 0 | 0 | 0 | 1 | 0 | 0 | 0 | 0 | 0 | 0 | 0 | 0 | 0 |
| 268/18 | O157:NT | NT | Clinical | 0 | 0 | 0 | 0 | 0 | 0 | 1 | 0 | 0 | 0 | 0 | 0 | 0 | 0 | 0 | 0 |
| 342/18 | O157:NM | NT | Clinical | 0 | 0 | 0 | 0 | 0 | 0 | 1 | 0 | 0 | 0 | 0 | 0 | 0 | 0 | 0 | 0 |
| 385/16 | O157:H7 | NT | Food | 0 | 0 | 1 | 1 | 0 | 0 | 1 | 1 | 0 | 1 | 0 | 0 | 1 | 0 | 1 | 0 |
| 566/18 | EAEC | NT | Clinical | 0 | 0 | 0 | 0 | 0 | 0 | 1 | 0 | 0 | 0 | 0 | 0 | 0 | 0 | 0 | 0 |
| 429/18 | EAEC | NT | Clinical | 0 | 0 | 0 | 0 | 0 | 0 | 1 | 0 | 0 | 0 | 0 | 0 | 0 | 0 | 0 | 0 |
| 126/18 | EAEC | NT | Clinical | 0 | 0 | 0 | 0 | 0 | 0 | 1 | 0 | 0 | 0 | 0 | 0 | 0 | 1 | 0 | 0 |
| 544/18 | EAEC | NT | Clinical | 0 | 0 | 0 | 0 | 0 | 0 | 1 | 0 | 0 | 0 | 0 | 0 | 0 | 0 | 0 | 0 |
| 434/18 | EAEC | NT | Clinical | 0 | 0 | 0 | 0 | 0 | 0 | 1 | 0 | 0 | 0 | 0 | 0 | 0 | 0 | 0 | 0 |
| 523/18 | EAEC | NT | Clinical | 0 | 0 | 0 | 0 | 0 | 0 | 1 | 0 | 0 | 0 | 0 | 0 | 0 | 0 | 0 | 0 |
| 451/17 | EAEC | NT | Clinical | 0 | 0 | 0 | 0 | 0 | 0 | 1 | 0 | 0 | 0 | 0 | 0 | 0 | 0 | 0 | 0 |
| 431/17 | EAEC | NT | Clinical | 0 | 0 | 0 | 0 | 0 | 0 | 1 | 0 | 0 | 0 | 0 | 0 | 0 | 0 | 0 | 0 |
| 417/17 | EAEC | NT | Clinical | 0 | 0 | 0 | 0 | 0 | 0 | 1 | 0 | 0 | 0 | 0 | 0 | 0 | 0 | 0 | 0 |
| 451/18 | O157:H7 | 2 | Clinical | 1 | 1 | 1 | 1 | 1 | 1 | 0 | 1 | 1 | 1 | 1 | 1 | 1 | 1 | 1 | 1 |
| 440/18 | O157:H7 | 2 | Clinical | 1 | 1 | 1 | 1 | 1 | 1 | 0 | 1 | 1 | 1 | 0 | 1 | 1 | 1 | 1 | 1 |
| 504/18 | O157:H7 | 2 | Clinical | 1 | 0 | 0 | 0 | 1 | 1 | 0 | 1 | 0 | 0 | 0 | 0 | 1 | 0 | 0 | 1 |
| 489/18 | O157:H7 | 2 | Food | 1 | 1 | 1 | 1 | 1 | 1 | 0 | 1 | 0 | 1 | 0 | 0 | 1 | 1 | 1 | 1 |
| 492/18 | O157:H7 | 2 | Clinical | 1 | 1 | 1 | 1 | 1 | 1 | 0 | 1 | 0 | 1 | 0 | 0 | 1 | 1 | 0 | 1 |
| 495/18 | O157:H7 | 2 | Clinical | 1 | 1 | 1 | 1 | 1 | 1 | 0 | 1 | 1 | 1 | 0 | 0 | 1 | 1 | 0 | 1 |
| 483/18 | O157:H7 | 2 | Clinical | 1 | 1 | 1 | 1 | 1 | 1 | 0 | 1 | 1 | 1 | 0 | 1 | 1 | 1 | 1 | 1 |
| 526/17 | O103:H3 | 1 | Clinical | 0 | 0 | 1 | 0 | 0 | 0 | 1 | 0 | 0 | 0 | 0 | 0 | 0 | 0 | 0 | 0 |
| 324/17 | O121:H19 | 2 | Clinical | 0 | 0 | 1 | 0 | 0 | 0 | 1 | 0 | 0 | 0 | 0 | 0 | 0 | 0 | 0 | 0 |
| 397/18 | O26:H11 | 1 | Clinical | 0 | 1 | 0 | 0 | 0 | 0 | 1 | 0 | 0 | 0 | 0 | 0 | 0 | 0 | 0 | 0 |
| 437/18 | O145:NM | 2 | Clinical | 0 | 1 | 0 | 0 | 0 | 0 | 1 | 0 | 0 | 0 | 0 | 0 | 0 | 0 | 0 | 0 |
| 488/18 | O145:NM | 2 | Clinical | 0 | 1 | 0 | 0 | 0 | 0 | 1 | 0 | 0 | 0 | 0 | 0 | 0 | 0 | 0 | 0 |
| 443/18 | O121:H19 | 2 | Clinical | 0 | 0 | 0 | 0 | 0 | 0 | 1 | 0 | 0 | 0 | 0 | 0 | 0 | 0 | 0 | 0 |
| 329/18 | ETEC | NT | Clinical | 0 | 0 | 0 | 0 | 0 | 0 | 0 | 0 | 0 | 0 | 0 | 0 | 0 | 0 | 0 | 0 |
| 330/18 | ETEC | NT | Clinical | 0 | 0 | 0 | 0 | 0 | 0 | 1 | 0 | 0 | 0 | 0 | 0 | 0 | 0 | 0 | 0 |
| 128/18 | EPEC | NT | Clinical | 0 | 0 | 0 | 0 | 0 | 0 | 1 | 0 | 0 | 0 | 0 | 0 | 0 | 0 | 0 | 0 |
| 221/18 | EPEC | NT | Clinical | 0 | 0 | 0 | 0 | 0 | 0 | 1 | 0 | 0 | 0 | 0 | 0 | 0 | 0 | 0 | 0 |
| 430/17 | EPEC | NT | Clinical | 0 | 0 | 0 | 0 | 0 | 0 | 1 | 0 | 0 | 0 | 0 | 0 | 0 | 0 | 0 | 0 |
| 611/17 | O157:H7 | 2 | Clinical | 1 | 1 | 1 | 1 | 1 | 1 | 0 | 1 | 1 | 1 | 0 | 0 | 1 | 1 | 1 | 1 |
| 744/17 | O157:H7 | 2 | Clinical | 1 | 1 | 1 | 1 | 1 | 1 | 0 | 1 | 1 | 1 | 0 | 0 | 1 | 1 | 1 | 1 |
| 736/17 | O157:H7 | 2 | Clinical | 1 | 1 | 1 | 1 | 1 | 1 | 0 | 1 | 1 | 1 | 0 | 0 | 1 | 1 | 0 | 1 |
| 732/17 | O157:H7 | 2 | Clinical | 1 | 1 | 1 | 1 | 1 | 1 | 0 | 1 | 1 | 0 | 0 | 0 | 1 | 1 | 0 | 1 |
| 715/17 | O157:H7 | 2 | Clinical | 1 | 1 | 1 | 1 | 1 | 1 | 0 | 1 | 1 | 1 | 0 | 0 | 1 | 1 | 0 | 1 |
| 718/17 | O157:H7 | 2 | Clinical | 1 | 1 | 1 | 1 | 1 | 1 | 0 | 1 | 1 | 1 | 1 | 0 | 1 | 1 | 0 | 1 |
| 594/17 | O157:H7 | 2 | Clinical | 1 | 1 | 1 | 1 | 1 | 1 | 0 | 1 | 0 | 0 | 0 | 0 | 1 | 1 | 0 | 1 |
| 17/19 | O157:H7 | 2 | Clinical | 1 | 1 | 1 | 1 | 1 | 1 | 0 | 1 | 0 | 0 | 0 | 1 | 1 | 0 | 1 | 1 |
| 18/19 | O157:H7 | 2 | Clinical | 1 | 1 | 0 | 1 | 1 | 1 | 0 | 1 | 0 | 0 | 1 | 1 | 1 | 0 | 1 | 1 |
| 34/19 | O157:H7 | 2 | Clinical | 1 | 1 | 1 | 1 | 1 | 1 | 0 | 1 | 0 | 0 | 0 | 1 | 1 | 0 | 0 | 1 |
| 19/19 | O157:H7 | 2 | Clinical | 1 | 1 | 0 | 1 | 1 | 1 | 0 | 1 | 0 | 0 | 0 | 1 | 1 | 1 | 1 | 1 |
| 38/19 | O157:H7 | 2 | Clinical | 1 | 1 | 1 | 1 | 1 | 1 | 0 | 1 | 0 | 1 | 0 | 1 | 1 | 0 | 1 | 1 |
| 39/19 | O157:H7 | 2 | Clinical | 1 | 1 | 0 | 1 | 1 | 1 | 0 | 1 | 1 | 1 | 0 | 0 | 0 | 0 | 1 | 1 |
| 704/18 | O157:H7 | 1 | Clinical | 1 | 1 | 1 | 1 | 1 | 1 | 0 | 0 | 0 | 0 | 0 | 0 | 1 | 0 | 1 | 1 |
| 731/18 | O157:H7 | 2 | Clinical | 1 | 1 | 1 | 1 | 1 | 1 | 0 | 0 | 0 | 0 | 0 | 0 | 1 | 0 | 1 | 1 |
| 770/18 | O157:H7 | 2 | Clinical | 1 | 1 | 1 | 1 | 1 | 1 | 0 | 1 | 0 | 0 | 0 | 0 | 1 | 0 | 1 | 1 |
| 772/18 | O157:H7 | 2 | Clinical | 1 | 1 | 1 | 1 | 1 | 1 | 0 | 1 | 0 | 0 | 0 | 0 | 1 | 0 | 0 | 1 |
| 771/18 | O157:H7 | 2 | Clinical | 1 | 1 | 1 | 1 | 1 | 1 | 0 | 0 | 0 | 0 | 0 | 0 | 1 | 0 | 1 | 1 |
| 727/18 | O157:H7 | 2 | Clinical | 1 | 0 | 1 | 1 | 1 | 1 | 0 | 1 | 0 | 1 | 1 | 1 | 1 | 1 | 1 | 1 |
| 726/18 | O157:H7 | 2 | Clinical | 1 | 0 | 1 | 1 | 1 | 1 | 0 | 0 | 0 | 0 | 0 | 1 | 1 | 1 | 0 | 1 |
| 687/18 | O157:H7 | 2 | Clinical | 1 | 0 | 1 | 1 | 1 | 1 | 0 | 0 | 0 | 0 | 0 | 1 | 1 | 1 | 0 | 1 |
| 660/18 | O157:H7 | 2 | Clinical | 1 | 0 | 1 | 1 | 1 | 1 | 0 | 0 | 1 | 0 | 0 | 0 | 0 | 0 | 0 | 1 |
| 602/18 | O157:H7 | 2 | Clinical | 1 | 1 | 1 | 1 | 1 | 1 | 0 | 1 | 1 | 1 | 0 | 0 | 1 | 1 | 0 | 1 |
| 714/18 | O157:H7 | 2 | Food | 1 | 1 | 0 | 0 | 0 | 0 | 0 | 0 | 0 | 0 | 0 | 0 | 0 | 0 | 0 | 0 |
| 657/18 | O157:H16 | NT | Clinical | 0 | 1 | 0 | 0 | 0 | 0 | 1 | 0 | 0 | 0 | 0 | 0 | 0 | 0 | 0 | 1 |
| 583/18 | O157:H7 | 2 | Clinical | 1 | 1 | 1 | 1 | 1 | 1 | 0 | 1 | 1 | 0 | 0 | 1 | 1 | 1 | 0 | 1 |
| 641/18 | O157:H7 | 2 | Clinical | 1 | 1 | 0 | 1 | 1 | 1 | 0 | 0 | 0 | 0 | 0 | 0 | 0 | 0 | 1 | 1 |
| 598/18 | O157:H7 | 2 | Clinical | 1 | 1 | 1 | 1 | 1 | 1 | 0 | 0 | 0 | 0 | 0 | 0 | 0 | 1 | 0 | 1 |
| 671/18 | O157:H7 | 2 | Clinical | 1 | 1 | 0 | 1 | 1 | 1 | 0 | 1 | 1 | 1 | 0 | 0 | 1 | 1 | 0 | 1 |
| 710/18 | O157:H7 | 2 | Clinical | 1 | 1 | 1 | 1 | 1 | 1 | 0 | 1 | 0 | 0 | 0 | 0 | 1 | 0 | 1 | 1 |
| 696/18 | O157:H7 | 2 | Clinical | 1 | 1 | 1 | 1 | 1 | 1 | 0 | 1 | 1 | 0 | 0 | 0 | 1 | 1 | 0 | 1 |
| 584/18 | O157:H7 | 2 | Clinical | 1 | 1 | 0 | 1 | 0 | 1 | 0 | 0 | 0 | 0 | 0 | 0 | 0 | 0 | 0 | 1 |
| 615/18 | O157:H7 | 2 | Food | 1 | 1 | 1 | 1 | 1 | 1 | 0 | 0 | 1 | 0 | 0 | 1 | 0 | 0 | 0 | 1 |
| 634/18 | O157:H7 | 2 | Food | 1 | 1 | 1 | 1 | 1 | 1 | 0 | 0 | 0 | 1 | 0 | 1 | 0 | 0 | 0 | 1 |
| 628/18 | O157:H7 | 2 | Food | 1 | 1 | 1 | 1 | 1 | 1 | 0 | 0 | 1 | 0 | 0 | 1 | 1 | 1 | 0 | 1 |
| 630/18 | O157:H7 | 2 | Food | 1 | 1 | 1 | 1 | 1 | 1 | 0 | 0 | 1 | 0 | 0 | 1 | 1 | 1 | 1 | 1 |
| 633/18 | O157:H7 | 2 | Food | 1 | 1 | 0 | 1 | 1 | 1 | 0 | 0 | 0 | 0 | 0 | 0 | 1 | 0 | 1 | 1 |
| 629/18 | O157:H7 | 2 | Food | 1 | 1 | 0 | 1 | 1 | 1 | 0 | 0 | 1 | 0 | 0 | 1 | 1 | 1 | 0 | 1 |
| 631/18 | O157:H7 | 2 | Food | 1 | 1 | 0 | 1 | 1 | 1 | 0 | 0 | 1 | 0 | 0 | 1 | 1 | 0 | 1 | 1 |
| 632/18 | O157:H7 | 2 | Food | 1 | 1 | 0 | 1 | 1 | 1 | 0 | 1 | 1 | 0 | 0 | 0 | 1 | 0 | 0 | 1 |
| 777/18 | O157:H7 | 2 | Clinical | 1 | 1 | 1 | 1 | 1 | 1 | 0 | 1 | 1 | 1 | 0 | 0 | 1 | 1 | 0 | 1 |
| 6 / 19 | O157:H7 | 2 | Clinical | 1 | 1 | 1 | 1 | 1 | 1 | 0 | 1 | 1 | 1 | 0 | 1 | 1 | 1 | 1 | 1 |
| 623/18 | O157:H7 | 2 | Clinical | 1 | 1 | 1 | 1 | 1 | 1 | 0 | 1 | 0 | 1 | 0 | 0 | 1 | 1 | 0 | 1 |
| 593/18 | O157:H7 | 2 | Clinical | 0 | 1 | 0 | 0 | 0 | 0 | 1 | 0 | 0 | 0 | 0 | 0 | 0 | 0 | 0 | 0 |
| 483/18 | EPEC | NT | Clinical | 0 | 0 | 0 | 0 | 0 | 0 | 1 | 0 | 0 | 0 | 0 | 0 | 0 | 0 | 0 | 0 |
| 518/17 | EPEC | NT | Clinical | 0 | 0 | 0 | 0 | 0 | 0 | 1 | 0 | 0 | 0 | 0 | 0 | 0 | 0 | 0 | 0 |
| 394/17 | EPEC | NT | Clinical | 0 | 0 | 0 | 0 | 0 | 0 | 1 | 0 | 0 | 0 | 0 | 0 | 0 | 0 | 0 | 0 |
| 166/18 | EAEC | NT | Clinical | 0 | 0 | 0 | 0 | 0 | 0 | 1 | 0 | 0 | 0 | 0 | 0 | 0 | 0 | 0 | 0 |
| 120/18 | EAEC | NT | Clinical | 0 | 0 | 0 | 0 | 0 | 0 | 1 | 0 | 0 | 0 | 0 | 0 | 0 | 0 | 0 | 0 |
| 295/18 | EAEC | NT | Clinical | 0 | 0 | 0 | 0 | 0 | 0 | 1 | 0 | 0 | 0 | 0 | 0 | 0 | 0 | 0 | 0 |
| 296/18 | EAEC | NT | Clinical | 0 | 0 | 0 | 0 | 0 | 0 | 1 | 0 | 0 | 0 | 0 | 0 | 0 | 0 | 0 | 0 |
| 300/18 | EAEC | NT | Clinical | 0 | 0 | 0 | 0 | 0 | 0 | 1 | 0 | 0 | 0 | 0 | 0 | 0 | 0 | 0 | 0 |
| 431/17 | EIEC | NT | Clinical | 0 | 0 | 0 | 0 | 0 | 0 | 1 | 0 | 0 | 0 | 0 | 0 | 0 | 0 | 0 | 0 |
| 417/17 | EIEC | NT | Clinical | 0 | 0 | 0 | 0 | 0 | 0 | 1 | 0 | 0 | 0 | 0 | 0 | 0 | 0 | 0 | 0 |
| 313/17 | EIEC | NT | Clinical | 0 | 0 | 0 | 0 | 0 | 0 | 1 | 0 | 0 | 0 | 0 | 0 | 0 | 0 | 0 | 0 |
| 309/17 | EIEC | NT | Clinical | 0 | 0 | 0 | 0 | 0 | 0 | 1 | 0 | 0 | 0 | 0 | 0 | 0 | 0 | 0 | 0 |
| 328/17 | EIEC | NT | Clinical | 0 | 0 | 0 | 0 | 0 | 0 | 1 | 0 | 0 | 0 | 0 | 0 | 0 | 0 | 0 | 0 |
| 361/17 | EIEC | NT | Clinical | 0 | 0 | 0 | 0 | 0 | 0 | 1 | 0 | 0 | 0 | 0 | 0 | 0 | 0 | 0 | 0 |
| 434/17 | EIEC | NT | Clinical | 0 | 0 | 0 | 0 | 0 | 0 | 1 | 0 | 0 | 0 | 0 | 0 | 0 | 0 | 0 | 0 |
| 493/18 | O157:H29 | NT | Clinical | 0 | 0 | 0 | 0 | 0 | 0 | 0 | 0 | 0 | 0 | 0 | 0 | 0 | 0 | 0 | 0 |
| 496/18 | O157:H29 | NT | Clinical | 0 | 0 | 0 | 0 | 0 | 0 | 1 | 0 | 0 | 0 | 0 | 0 | 0 | 0 | 0 | 0 |
| 382/18 | E coli | S/FV | Clinical | 0 | 0 | 0 | 0 | 0 | 0 | 1 | 0 | 0 | 0 | 0 | 0 | 0 | 0 | 0 | 0 |
| 149/18 | E coli | S/FV | Clinical | 0 | 0 | 0 | 0 | 0 | 0 | 1 | 0 | 0 | 0 | 0 | 0 | 0 | 0 | 0 | 0 |
| 342/18 | O157:NM | NT | Clinical | 0 | 0 | 0 | 0 | 0 | 0 | 1 | 0 | 0 | 0 | 0 | 0 | 0 | 0 | 0 | 0 |
